# The evolutionary origins and diversity of the neuromuscular system of paired appendages in batoids

**DOI:** 10.1101/644062

**Authors:** Natalie Turner, Deimante Mikalauskaite, Krista Barone, Kathleen Flaherty, Gayani Senevirathne, Noritaka Adachi, Neil H Shubin, Tetsuya Nakamura

## Abstract

Appendage patterning and evolution have been active areas of inquiry for the past two centuries. While most work has centered on the skeleton, particularly that of amniotes, the evolutionary origins and molecular underpinnings of the neuromuscular diversity of fish appendages have remained enigmatic. The fundamental pattern of segmentation in amniotes, for example, is that all muscle precursors and spinal nerves enter either the paired appendages or body wall at the same spinal level. The condition in finned vertebrates is not understood. To address this gap in knowledge, we investigated the development of muscles and nerves in unpaired and paired fins of skates and compared them to those of chain catsharks. During skate and shark embryogenesis, cell populations of muscle precursors and associated spinal nerves at the same axial level contribute to both appendages and body wall, perhaps representing an ancestral condition of gnathostome appendicular neuromuscular systems. Remarkably in skates, this neuromuscular bifurcation as well as colinear *Hox* expression extend posteriorly to pattern a broad paired fin domain. In addition, we identified migratory muscle precursors (MMPs), which are known to develop into paired appendage muscles with *Pax3* and *Lbx1* gene expression, in the dorsal fins of skates. Our results suggest that muscles of paired fins have evolved via redeployment of the genetic program of MMPs that were already involved in dorsal fin development. Appendicular neuromuscular systems most likely have emerged as side branches of body wall neuromusculature and have been modified to adapt to distinct aquatic and terrestrial habitats.

## Introduction

Emergence and diversification of paired appendages are central to vertebrate evolution (1). During evolution of paired appendages, skeletons, muscles and nerves have been assembled to support appendage movement, but the evolutionary trajectories and diversity of underlying genetic mechanisms of appendicular neuromuscular patterning remain largely unknown (2). Cartilaginous fishes, consisting of chimaeras, sharks, skates, and rays, hold prominent phylogenetic positions in vertebrate evolution, representing primitive conditions of paired appendages (3, 4). In addition to their significance in evolutionary studies, cartilaginous fishes exhibit remarkably diverse paired fins. For example, skates, rays (batoids), and angel sharks, have evolved extraordinarily broad paired fins (5). To generate power for forward propulsion, batoids primarily rely on undulatory movement of wide pectoral fins (6). This motion is achieved by a unique arrangement of the skeleton, muscles, and nerves (7). Despite their phylogenetically significant position, functional variety, and evolutionary diversity, the developmental processes and mechanisms of appendicular neuromusculature have not been investigated in diverse cartilaginous fish.

In amniotes, hypaxial muscle precursors develop ventral and appendicular musculature to emerge from the ventrolateral dermomyotome of somites during embryogenesis (8). These muscle precursors are categorized into subpopulations based on their positions along the anteroposterior axis (9). In limb segments, hypaxial muscle precursors delaminate from the dermomyotome and migrate into the limb bud as migratory muscle precursors (MMPs). At interlimb levels, the ventral lip of the dermomyotome directly extends into the body wall forming body wall muscles (9). Intriguingly, analysis of developmental processes of appendage and body wall muscles in chondrichthyans and other fish have identified differences to the condition in amniotes (10, 11). Whereas MMPs migrate and differentiate into appendage muscles in amniotes, appendage muscles of chondrichthyans have been argued to be derived from a direct extension of the ventrolateral dermomyotome into the pectoral fin (12, 13). It has also recently been shown that catsharks have delaminated MMPs entering into the pectoral fin, a seemingly comparable mechanism with that of amniotes (10). Furthermore in amniotes, hypaxial muscle precursors migrate exclusively either into the body wall or paired appendages (9). In catsharks, hypaxial muscle precursors emigrate into both (2, 10). Lacking comparative data, particularly from cartilaginous fish, we currently do not know the phylogenetic polarity or significance of these changes to developmental patterning.

Genetic underpinnings of appendage and body wall muscle development have been revealed in mouse and chick embryos. At limb segments, MMPs express the homeodomain transcription factor *Lbx1 (ladybird homeobox 1).* Loss of *Lbx1* results in a lack of appendage muscles and MMP migration (14–16), indicating that *Lbx1* function is critical for the migratory ability of MMPs. In contrast, direct extension of the ventrolateral dermomyotome at the interlimb level takes place without expression of *Lbx1* (9, 17). A previous study suggested that position-dependent expression of *Lbx1* in hypaxial muscle precursors along the anteroposterior axis is determined by combinations of *Hox* genes in mice (18).

In vertebrates, concomitant with musculature development, motoneurons (MNs) originate from the ventral neural tube and extend to and innervate their target muscles. Coincident with *Hox* expression along the anteroposterior axis, MNs differentiate into lateral motor column neurons (LMC) that innervate paired appendage muscles at the brachial and lumbar levels, hypaxial motor column neurons (HMC) that innervate hypaxial muscles at the thoracic and sacral levels, or medial motor column neurons (MMC) that regulate epaxial muscles (19). At brachial and lumber levels, *Hox6* and *Hox10* genes induce high expression level of *FoxP1*, a transcription factor that promotes LMC differentiation via direct binding to its regulatory regions (20). In contrast, at the thoracic level, *Hoxc9* represses *FoxP1* expression and induces development of the HMC (21). Intriguingly, skates (*Leucoraja erinacea*) and catsharks (*Scyliorhinus canicular*) lost the *HoxC* cluster during evolution (22, 23). Loss of the *HoxC* cluster results in high *FoxP1* expression, and the through development of the LMC between brachial and lumbar domains in skates (24). Unexpectedly, a previous study identified a minor population of MNs that is similar to the HMC at pectoral and pelvic domains with LMC in the neural tube of skates (24), implying that skates retain unique patterns of innervation at the brachial level compared with other finned vertebrates. However, developmental patterns of peripheral nerves in skates and other rays have not yet been sufficiently investigated.

To understand how batoid neuromuscular systems arose, we investigated their developmental patterns in skates (*Leucoraja erinacea*) and compared them with chain catsharks (*Scyliorhinus retifer*) using whole-mount *in situ* hybridization and antibody staining. These data provide molecular insights into the evolutionary mechanisms behind the generation of diverse neuromuscular systems as well as their evolutionary origins.

## Results

### Dual contribution of muscles into the body wall and pectoral fin

To understand the developmental mechanisms of cephalic and appendicular muscles in batoids, we performed whole-mount *in situ* hybridization for *Pax3* and *Lbx1* mRNA in skate embryos. *Pax3* was expressed in myoblasts of the pectoral fin and body wall (Fig. 1A,B). Subsequent sectioning of stained embryos confirmed *Pax3* expression in the dermomyotome, myoblasts of the pectoral fin, and ventrally extended body wall myoblasts (Supplementary Fig. 1). This dual contribution pattern of *Pax3* expression in the pectoral fins and body wall is similar to that of shark embryos (10), although it was extended further posteriorly in skate fins compared to shark fins. Similarly, *Lbx1* expression was observed in myoblasts of the pectoral fin, but not in the body wall, unlike *Pax3* expression (Fig.1C). This suggests that the myoblast population derived from the dermomyotome separates into fin and body wall muscles with and without *Lbx1* expression, as MMPs and muscle progenitor cells, respectively.

**Figure 1.**
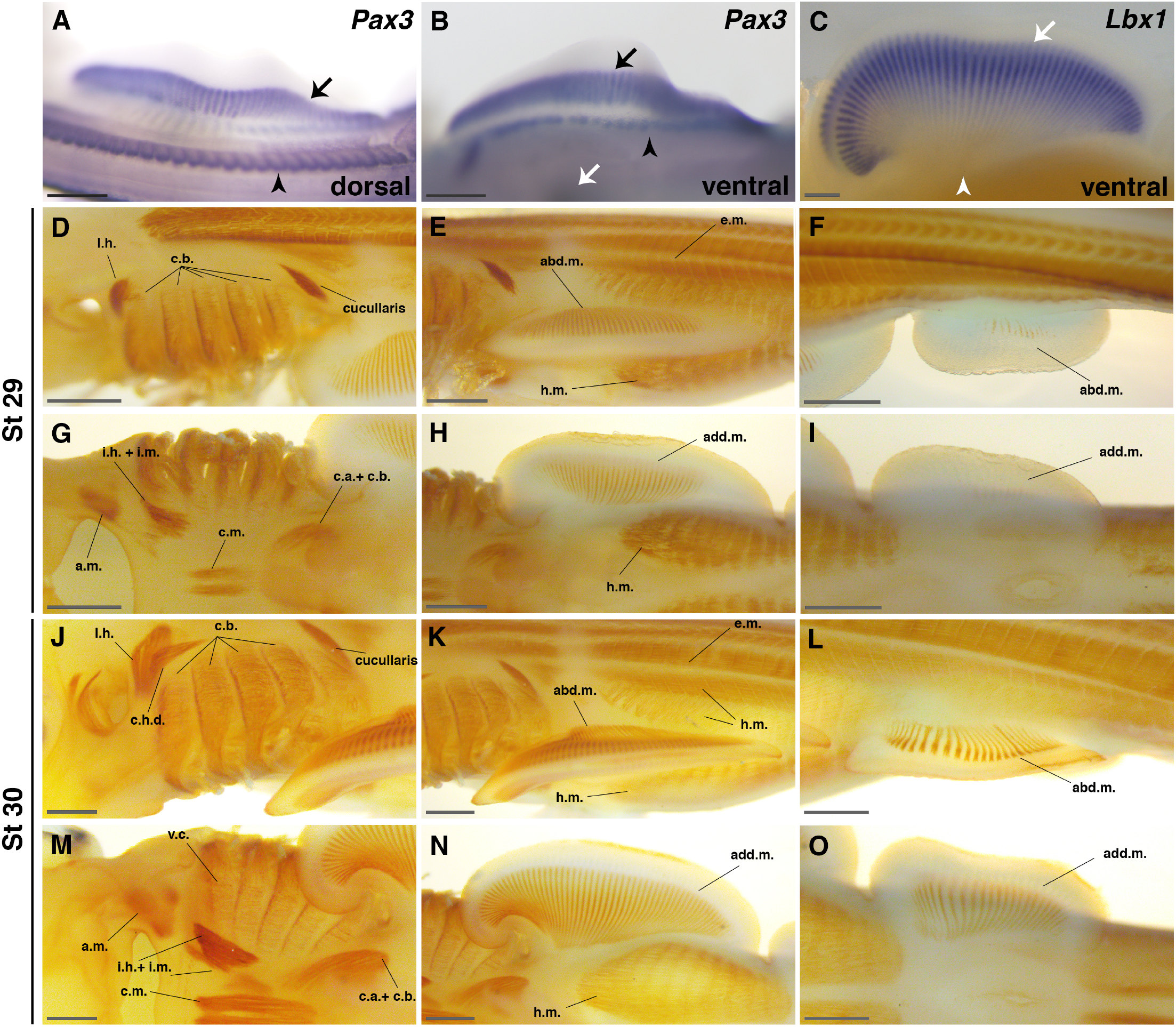
Musculature development in paired fins of skate. (**A-C**) Whole-mount *in situ* hybridization of *Pax3* and *Lbx1*. (**A, B**) Dorsal and ventral views of the expression patterns of *Pax3* in the pectoral fin. *Pax3* is expressed in both muscles of the pectoral fin (arrow) and dermomyotome (arrowhead) (**A**), or the pectoral fin (arrow) and body wall muscles (arrowhead) (**B**). White arrow points to the umbilical cord (**B**). (**C**) Ventral view of *Lbx1* expression pattern in the pectoral fin. *Lbx1* is expressed only in the pectoral fin (arrow) and not in the body wall muscles (arrowhead). (**D-O**) Immunostaining of skate embryos by myosin heavy chain antibody at stage 29 (**D-I**) or stage 30 (**J-O**) in lateral (**D, E, F, J, K, and L**) or ventral (**G, H, I, M, N, and O**) view. At stage 29, the constrictor branchialis, cucullaris and other cephalic muscles develop (**D, G**). Abductor and adductor muscles start to develop in the pectoral (**E, H**) and pelvic fins (**F, I**), yet they do not fully develop distally. Note that the abductor and adductor muscles of the pectoral fin and hypaxial muscles of the body wall develop at the same axial level (**E, H**). At stage 30, cephalic muscles are more developed compared to stage 29. Cucullaris extends dorsal to branchial arches (**J**). Interhyoideus and coracomandibularis are clearly identified (**M**). Abductor and adductor muscles in the pectoral fins (**K, N**) and pelvic fins (**L, O**) develop towards the distal direction. Abbreviations; abd.m.; abductor muscles, add.m.; adductor muscles, a.m.; adductor mandibulae, c.a. + c.b.; a complex of coraco arcualis and coracobranchialis, c.h.d.; constrictor hyoideus dorsalis, c.m.; coracomandibularis, e.m.; epaxial muscles, h.m.; hypaxial muscles, i.h. + i.m.; a complex of interhyoideus and intermandibularis, l.h.; levator hyomandibulae. All scale bars are 0.5 mm.

We next performed whole-mount immunostaining of skate embryos using myosin heavy chain antibodies. Basic components of cephalic and appendicular muscles were observed at stages 29 and 30 of embryonic development (Fig. 1D–O). The constrictor branchialis, which is necessary for gill movement, developed in the pharyngeal arches (Fig. 1D). The cucullaris, which articulates the skull and pectoral girdle, was formed at dorsal to the pharyngeal arches and lateral to epaxial muscles (Fig. 1D,E). Hypobranchial muscles developed at the ventral gill region (Fig. 1G). In the pectoral fin, adductor and abductor muscles were observed at the dorsal and ventral sides, respectively (Fig. 1E,H). Hypaxial body wall muscles developed posteriorly from the base of the umbilical cord and backward, consistent with *Pax3* expression patterns (Fig. 1E, H, K, N). This embryonic pattern is distinct from tetrapod musculature, in which the early development of appendage and body wall muscles occurs exclusively along the anteroposterior axis. Adductor and abductor muscles were formed in the pelvic fin as well, but staining of myosin heavy chain was weaker than those of the pectoral fin at stage 29 (Fig. 1F,I). At stage 30, abductor and adductor muscle staining was stronger and extended to the distal tip of pectoral and pelvic fins compared to stage 29 (Fig. 1K,N,L,O).

### Branching of spinal nerves into the body wall and pectoral fins

A previous study reported that the localization of LMC neurons expands posteriorly in the neural tube of skates (24). However, the peripheral innervation pattern of batoid spinal nerves during embryogenesis has not been studied in detail. To investigate developmental patterns of skate nerves, we used a 3A10 antibody that recognizes neurofilament-associated proteins. In contrast to the contribution of Sp nerves 5-8 into the pectoral appendage in mice (25), Sp nerves 1–32 innervated into the pectoral fin muscles in skate (Fig. 2A). Particularly, Sp nerves 1–10 developed the brachial plexus and innervated anterior pectoral fin muscles (Fig. 2A–C). In the posterior part of the pectoral fin, Sp nerves bifurcated at the base of the pectoral fin; one branch innervated body wall muscles (ventral branch of Sp nerves) and the other branch entered pectoral fin muscles at the same axial level (pectoral nerve) (Fig. 2D). After entering into the pectoral fin, pectoral nerves branched into dorsal and ventral domains for adductor and abductor muscles, respectively. In contrast to the previous study, however, we did not confirm a contribution of Oc nerves into pectoral fin muscles. Similar to pectoral nerves, Sp nerves 33–48 branched and innervated into the pelvic fins and body wall (Fig. 2A,E).

**Figure 2.**
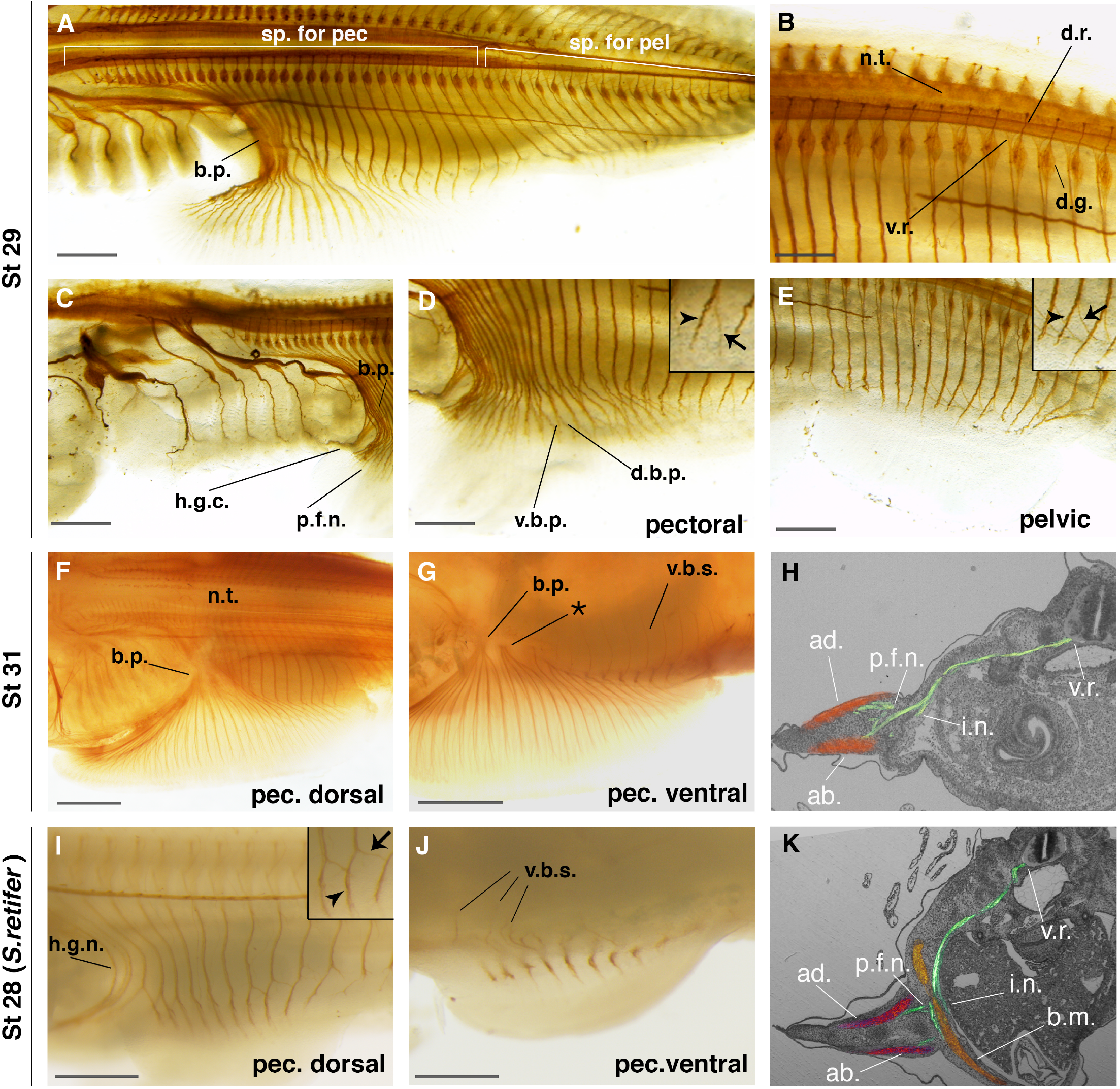
Developmental pattern of nerves in chondrichthyan paired fins. Immunostaining of nerves in developing embryos of skate (*L. erinacea)* and shark (*S. retifer)* with 3A10 antibody. (**A-E**) Skate embryos at stage 29. (**A**) Dorsal view shows the innervation patterns for the pectoral and pelvic fins. The brachial plexus consists of Sp nerves. (**B**) Dorsal (sensory) and ventral (motor) root of spinal nerves for paired appendages. (**C**) Lateral view of the branchial arch domain. Occipital nerves do not contribute to the brachial plexus; only spinal nerves form the brachial plexus. The hypoglossal nerve and the pectoral fin nerves were observed. (**D**) Dorsolateral view of the pectoral fin region. Spinal nerves first branch into the pectoral fin laterally (arrowhead) and the body wall muscle ventrally (arrow) (shown in inset). Then, a second branching event occurred and the pectoral fin nerves innervate the dorsal and ventral muscles of the pectoral fin. (**E**) Dorsolateral view of the pelvic fin region. Inset shows the branching event where the spinal nerves split and innervate the pelvic fin (arrowhead) and body wall (arrow) muscles. (**F-G**) Skate embryos at stage 31. (**F**) Dorsal view of the pectoral fin region. Sp nerves directly innervate the posterior pectoral fin without forming the brachial plexus. (**G**) Ventral view of the pectoral fin region showing the brachial plexus and a second plexus-like structure (*) innervating the pectoral fin. (**H**) 3D reconstruction of neuromuscular systems in skate embryos at stage 29. The ventral **roots** of Sp nerves (green) **exit** from the ventral neural tube and innervate into dorsal and ventral pectoral fin (orange) as well as body wall muscles. (**I-J**) Shark embryos at stage 28. (**I**) Dorsal view of the pectoral region. Inset shows the branching event as the Sp nerves split and innervate the pectoral fin (arrowhead) and the body wall (arrow) muscles. Distally, the pectoral fin nerves branch dorsoventrally. (**J**) Ventral view of the pectoral fin. The v.b.s. is observed in the body wall. (**K**) 3D reconstruction of neuromuscular systems in shark embryos at stage 29. The ventral roots of Sp nerves (green) innervate into dorsal and ventral pectoral fin (red) as well as body wall muscles (yellow). Abbreviations; a.b.; abductor muscle of the pectoral fin, a.d.; adductor muscle of the pectoral fin, b.m.; body wall muscle, b.p.; brachial plexus, d.b.p.; dorsal branch of the pectoral nerve, d.g.; dorsal root ganglion, d.r.; dorsal root, h.g.n.; hypoglossal nerve, v.b.; ventral branch of spinal nerves, n.t.; neural tube, p.f.n.; pectoral fin nerves, sp.; spinal nerves, v.b.p.; ventral branch of the pectoral nerve, v.b.s.; ventral branch of the spinal nerve, v.r.; ventral root. All scale bars are 1 mm.

At stage 31, Sp nerves established a brachial plexus-like structure in addition to the brachial plexus of Sp nerves 1–10 at the middle of the pectoral fin. Dorsal and ventral fin muscles were also innervated by Sp nerves at this stage (Fig. 2F,G). The plexus-like nerve bundle may be indispensable to innervate into the pectoral fin through the pectoral girdle. The posterior pectoral fin was innervated by Sp nerves that branched at the base of the fins and extended into the body wall and pectoral fin without forming plexus structures.

### Conserved developmental pattern of muscles and nerves in chondrichthyans

Our results showed that myoblast population and spinal nerves contribute into both the pectoral fin and body wall at the same vertebrae level during skate embryogenesis (Fig. 1 and Fig. 2). A previous study identified the similar migratory pattern of myoblasts in catsharks (10). To further compare developmental patterns of pectoral fin muscles and nerves of skates with those of sharks, we used myosin heavy chain antibodies to stain developing muscles and nerves in chain catshark embryos (*Scyliorhinus retifer*). At stage 30, staining confirmed abductor and adductor muscles in the pectoral fin (Supplementary Fig. 2). In the trunk, hypaxial body wall muscles developed at the same axial level as the pectoral fin muscles (Supplementary Fig. 2). In addition, Sp nerves 7–16 bifurcated into the body wall and pectoral fin without forming a plexus structure at stage 28 (Fig. 2I,J).

To confirm peripheral structures of Sp nerves, we stained serial sections of skate and shark embryos from stages 29 to 32 with hematoxylin and eosin solutions. After photographing serial sections, we reconstructed the original 3D morphology of muscles and nerves by using Amira software (Materials and Methods). 3D reconstruction of skate and shark embryos showed that Sp nerves that originated from the ventral root of neural tube innervated both pectoral fin and body wall muscles peripherally, indicating that they contain motor nerves that regulate movement of body wall and pectoral fin muscles (Fig. 2H and K). Collectively, these results suggest that skates and sharks retain comparable developmental patterns of neuromuscular systems, in which muscles and nerves contribute into both the paired appendages and body wall at the same vertebrae level, and batoids have posteriorly extended this dual contribution compared with sharks, supporting movement of their exceptionally wide fins.

### Dynamic rearrangements of *Hox* expression patterns in skates

Previous studies have revealed evolutionary diversity of MNs for paired appendages of elephant sharks, skates, and zebrafish (24, 26). However, the molecular mechanisms that coordinate evolution of appendage muscles, nerves, and skeletons remains unknown. Particularly in skates, development of the pectoral fin initiates from a strikingly wide fin bud that derives from the lateral plate mesoderm (LPM) (27, 28). Subsequently, muscles and nerves enter the enlarged pectoral fin bud from wider domains of the paraxial mesoderm (PAM) and neural tube along the anteroposterior axis compared with shark embryos. This observation led us to hypothesize that vertebrates have mechanisms that coordinate development of muscles, nerves, and skeletons to support appendage movement.

To understand the genetic mechanisms involved in co-evolution of muscles, nerves, and skeletons in skate, we investigated gene expression patterns of *Fgf8 (fibroblast growth factor 8)*, *Wnt3*, *Cyp26a1 (cytochrome P450 26a1)*, and *Cdx2*, which provide anteroposterior positional information during gastrulation and specify prospective limb regions (29–31). Expression patterns of these genes at stage 23 did not differ remarkably from those of other vertebrates at the comparable stage (Fig. 3 A–D). To further explore the genetic mechanisms for co-evolution, we tested expression patterns of *Hox* genes which provide positional information along the anteroposterior axis during gastrulation (32, 33) as the downstream targets of *Fgf8, Wnt3*, *Cyp26a1* and *Cdx2* in skates. Consistent with *Hox* expression patterns of catsharks (34), *Hoxa2*, *Hoxa3*, *Hoxa4, Hoxd1, Hoxd3 and Hoxd4* were expressed from the rhombomeres to the caudal tip of neural tube (Fig. 3E,F,G,M,N,O,Q). However, comparison of the anterior limit of *Hoxa9*, *Hoxa10*, *Hoxa11*, and *Hoxd8* expression showed that their expression domains shifted posteriorly in the neural tube of skates compared to catsharks (34) (Fig. 3I,J,K,P,Q, Materials and Methods). In the PAM of skate embryos, expression domains of *Hoxa9*, *Hoxa10*, and *Hoxa11* also shifted posteriorly from that of sharks (Fig. 3I,J,K,Q). Furthermore, in skate LPM, *Hoxa4*, *Hoxd4*, and *Hoxa5*, which are capable of inducing *Tbx5* expression via direct binding to regulatory regions in mice (35), exhibited broader expression domains than in sharks and other tetrapods (Fig. 3G,H,O,Q). These results show that dynamic rearrangements of *Hox* expression patterns have occurred in the neural tube, PAM, and LPM of skates.

**Figure 3.**
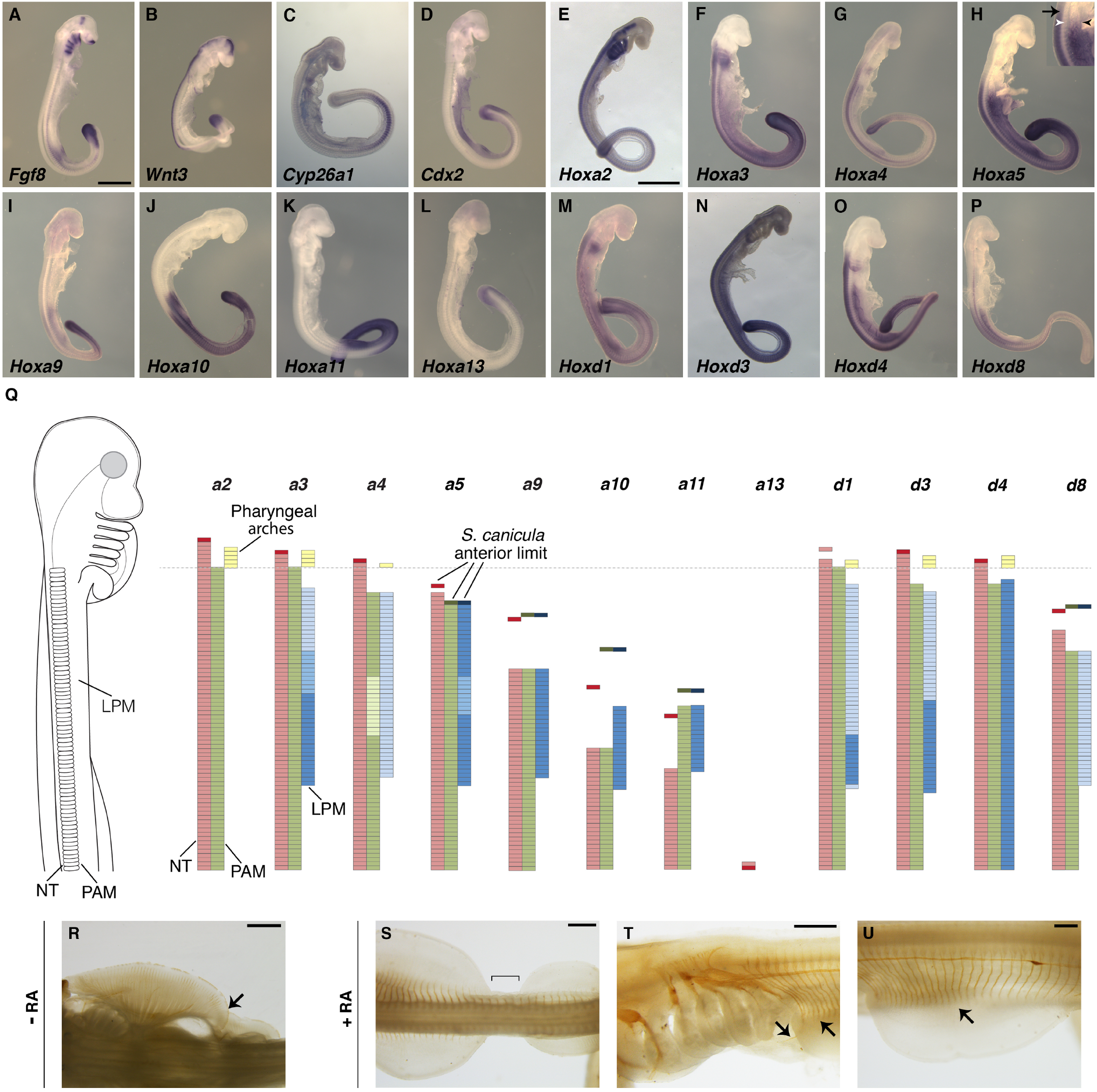
Expression pattern of *Hox* genes during skate gastrulation. (**A-P**) Whole mount *in situ* hybridization of *Fgf8*, *Wnt3*, *Cyp26a1*, *Cdx2*, and *Hox* groups A and D in *L. erinacea* at stage 23. Note that *Hox* genes show colinear expression along the anteroposterior axis. In inset H, the black arrow, the white arrowhead, and the black arrowhead point to the anterior limits of the expression in the neural tube, PAM, and LPM, respectively. All scale bars are 1 mm. The photos are scaled except for E, G, N, and P, which are scaled with each other. (**Q**) Schematic summary of *Hox* expression patterns in *L. erinacea* in the neural tube (red), pharyngeal arches (yellow), paraxial mesoderm (green), and lateral plate mesoderm (blue). Expression levels of *Hox* genes are indicated by color darkness. Darker bars show the anterior limit of expression of each *Hox* gene in *S. retifer* embryos (34), indicating that expression patterns of *Hoxa9*, *a10, a11*, and *d8* have shifted posteriorly in *L. erinacea.* Anterior limits of each gene in the neural tube (NT), lateral plate mesoderm (LPM), and paraxial mesoderm (PAM) were determined by extending the anterior border of the adjacent somite (Material and Methods). (**R**) Innervation staining of an RA-*L. erinacea* embryo. Note the overlap of the pectoral and pelvic fins (arrow). (**S-U**) Innervation staining of *L. erinacea* embryos cultured with retinoic acid, resulting in narrower pectoral fin size than RA-embryos and creating a thoracic region between the pectoral and pelvic fins (bracket) (**S**). (**T**) The brachial plexus did not develop in embryos treated by RA, although nerves innervate hypobranchial and pectoral muscles (arrows). (**U**) Spinal nerves exclusively innervate the pectoral fin in embryos treated by retinoic acid (arrow). All scale bars are 1 mm.

To test the effects of rearrangements of *Hox* expression patterns on the development of pectoral fins in skates, we cultured skate embryos with retinoic acid (RA), which alters *Hox* expression patterns (36), from stage 23 to 30 (Fig. 3 R-U). Pectoral fins of embryos cultured with RA were narrower than control embryos along the anteroposterior axis (9/10 embryos; Fig. 3R,S and Supplementary Fig. 3), creating a thoracic domain between pectoral and pelvic fins. RA–treated embryos lost the brachial plexus (Fig. 3T) and the branching pattern of pectoral nerves— Sp nerves directly innervated the pectoral fin from the neural tube (Fig.3U). These results suggest that *Hox* expression is responsible for regulating the width of skate fins, plexus formation, and innervation patterns.

### MMPs in dorsal fin development

While developmental processes of pectoral muscles and nerves have been investigated in multiple fish taxa (2, 37), evolutionary origins of the developmental programs for neuromuscular systems in paired appendages remain unexplored. Therefore, we investigated *Pax3* expression that represents muscle precursor cells by whole-mount *in situ* hybridization. *Pax3* is expressed weakly at the proximal base of 1^st^ and 2^nd^ dorsal fins at stage 29 (Fig. 4A), indicating that muscle precursor cells are present in developing dorsal fins. Then, to test whether these *Pax3*–positive cells are MMPs, we investigated *Lbx1* expression in skate dorsal fins. At stage 29, *Lbx1* begins to be expressed in the proximal domains of first and second dorsal fins, which is similar to *Pax3* expression (Fig. 4B). At stage 30, *Lbx1* expression extended to the distal part of dorsal fins and was undetectable at stage 31 (Fig. 4C). Sectioning of embryos after whole-mount *in situ* hybridization showed that *Lbx1*-expressing cells resided beneath the epidermis of dorsal fins (Fig. 4D). Furthermore, actin staining by phalloidin showed that MMPs accumulate actin internally, which is a signature of differentiating muscle precursor cells (Fig. 4E,F). These results demonstrate that the dorsal fin muscles of skates originate from MMPs.

**Figure 4.**
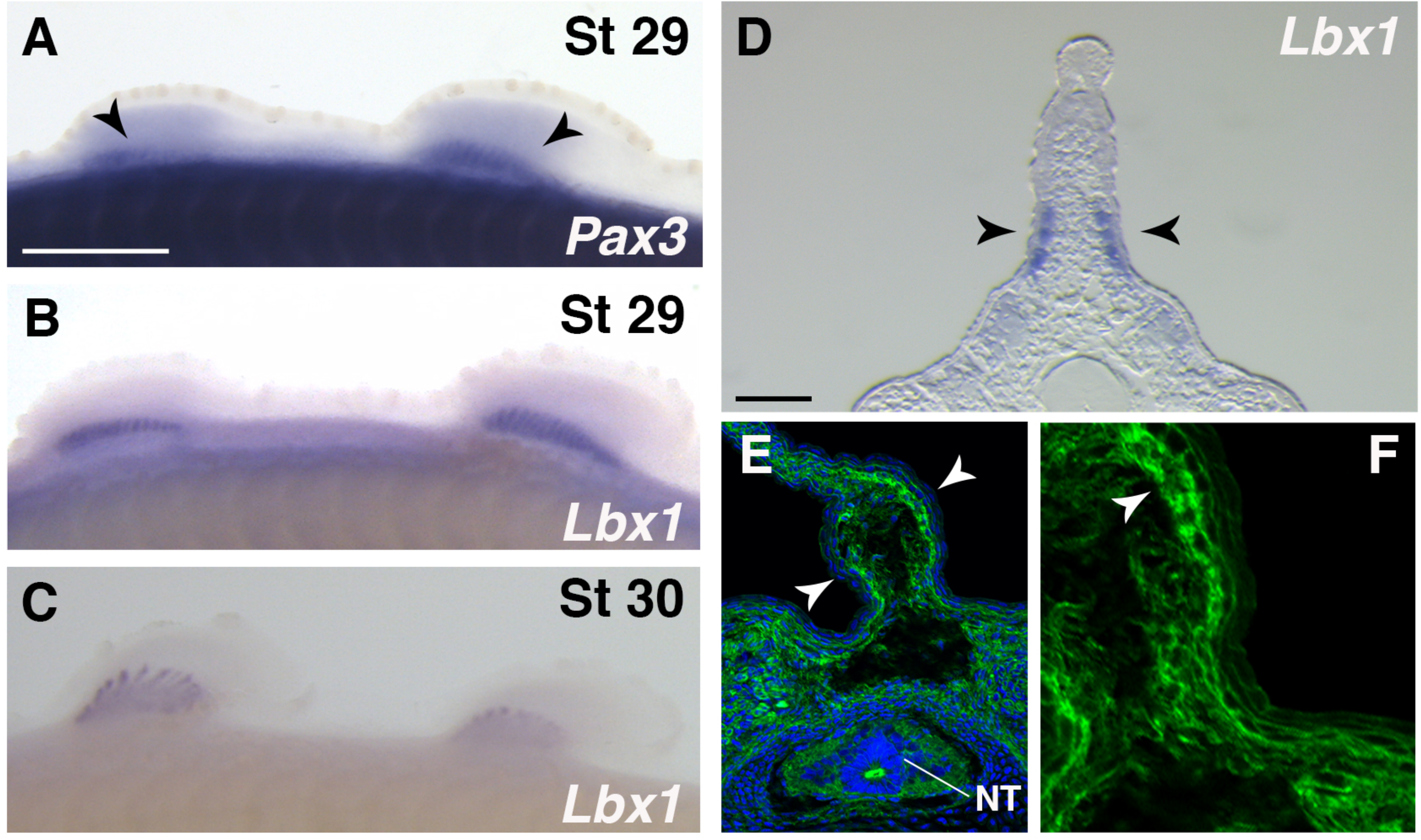
Migratory muscle precursor cells in the dorsal fins. (**A**) *Pax3* expression in the first and second dorsal fins at stage 29. The expression is confirmed at the base of the dorsal fins (arrows). (**B, C**) *Lbx1* expression in the dorsal fins at stage 29 (**B**) and 30 (**C**). *Lbx1* is highly expressed in the muscle precursor cells in the first and second dorsal fins at stage 29 and the expression extends in the distal direction at stage 30. (**D**) Section of the embryo stained by whole-mount *in situ* hybridization of *Lbx1*. Arrowheads indicate expression of *Lbx1*. (**E, F**) Section staining of a dorsal fin by phalloidin (Actin; green) and DAPI (nucleus; blue). Migratory muscle precursor cells are observed right under the epidermal tissues. F is a magnified image of E without overlay of DAPI. All scale bars are 1 mm.

## Discussion

Comparison of skates and chain catsharks highlights a conserved developmental pattern of neuromuscular systems – the dual contribution into the body wall and paired appendages at the same vertebrae level. Intriguingly, the dual contribution of hypaxial muscle precursors into the body wall and pectoral fins has been previously reported in species of catsharks (10). Furthermore, the branching pattern of spinal nerves into the pectoral fin and body wall was also described in the anatomical study of dogfish (*Squalus acanthias)*, lungfish (*Propterus dolloi*), and teleosts (26, 38–40). The cumulative knowledge of appendage neuromuscular systems in diverse taxa implies that their extension into both the body wall and paired appendages at the same vertebrae level, which is conserved in chondrichthyans, actinopterygians, and sarcopterygians, and absent in agnathans, is a synapomorphy for gnathostome appendages (Fig. 5).

**Figure 5.**
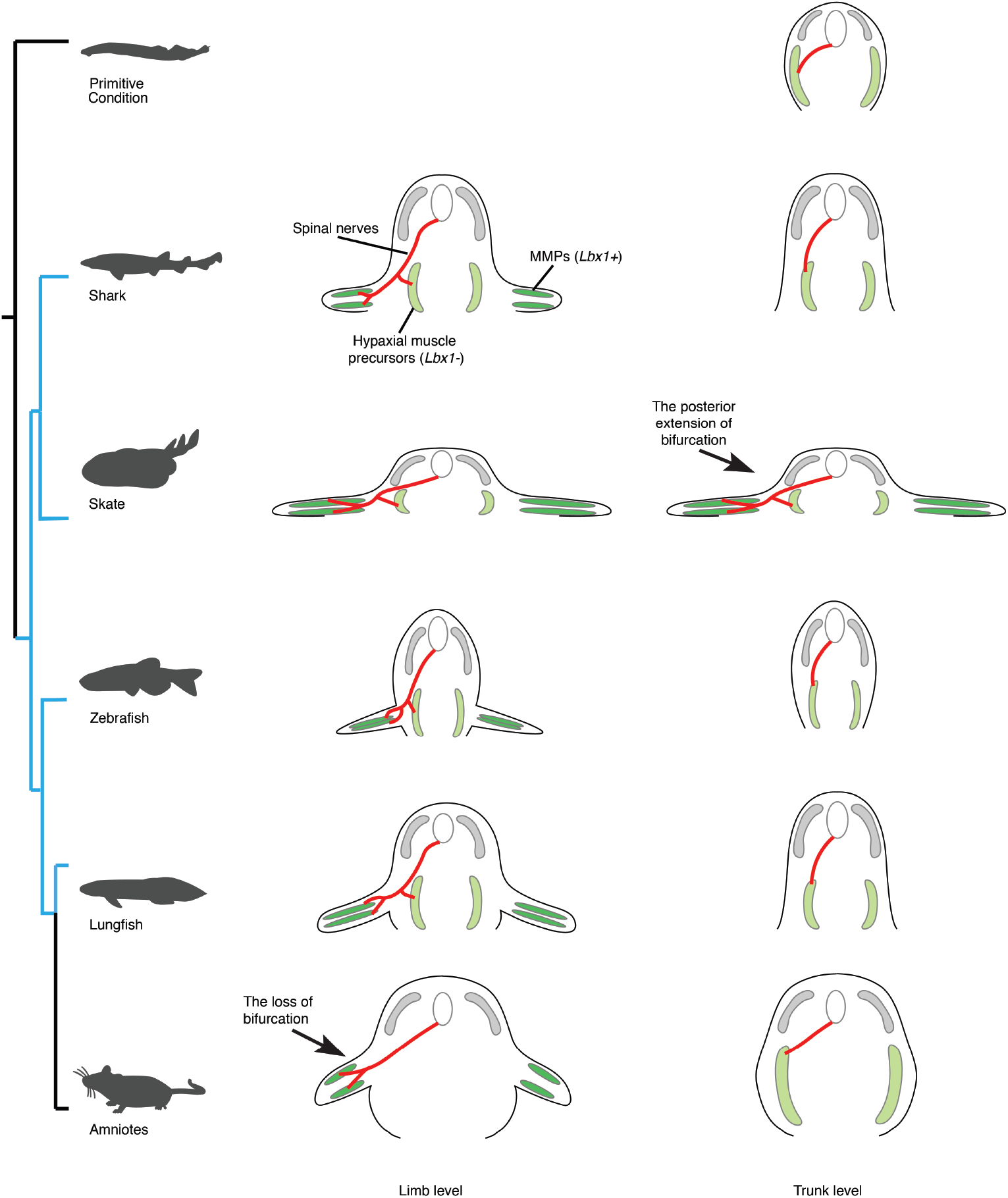
Summary and hypothesis of neuromuscular evolution in appendages. Summary of neuromuscular evolution in appendages. In skates, sharks, zebrafish, and lungfish, muscles and nerves branch into the body wall and paired appendages at the same axial level (Blue color in the tree represents animals that possess the dual contribution of neuromuscular systems). Amniotes lost this branching pattern, and neuromuscular components exclusively contribute to either body wall or paired appendages. In skates, the dual contribution of neuromuscular components is extended posteriorly to support undulatory swimming.

In contrast to ancestral dual contribution of the neuromuscular systems in fish, tetrapods do not possess similar patterns at the brachial segments (Fig.5). This loss may be related to appendage narrowing: the narrowing width of pectoral appendages during the fish-to-tetrapod transition (41, 42), may have been associated with the loss of this feature. In mammals, rostral intercostal nerves (the 2nd and 3rd in humans) have the intercostobrachial branches, and the subclavius nerve branches from the brachial plexus, both of which extend into the pectoral and body wall domains (43). These nerves may be a remnant of the branching nervous system from the primitive condition of vertebrates, although comparative analysis of their developmental patterns as well as underlying molecular mechanisms across diverse taxa is critical to conclude.

How has the pattern of dual contribution of neuromuscular systems been modified or lost during the evolution of limbed vertebrates? In chicken embryos, *Hoxa4* is expressed in the LPM lateral to somites 3–7 and *Hoxa5* is expressed in the LPM adjacent to somites 4–10, both which are capable of inducing *Tbx5* expression (35). In skates, these *Hox* genes are broadly expressed in the LPM lateral to somites 7–50, which is significantly wider than other vertebrates (Fig. 3Q). While LPM expression domains of *Hox* paralogous groups 4 and 5 in sharks are not precisely described, these dynamic changes of *Hox* expression patterns in the LPM are most likely involved in evolving wide pectoral fins. *Hox* expression is also indispensable for hindlimb development. Particularly, *Hoxa9*–*11* genes likely determine hindlimb position in the LPM and vertebrae identity in the PAM (44, 45). In skate, *Hoxa9*, *Hoxa10*, *Hoxa11, and Hoxd8* are shifted caudally compared with sharks in the neural tube, LPM, and PAM (Fig. 3Q). The posterior shift of these *Hox* genes probably caused extensive and concomitant remodeling of muscles, nerves, and skeletons in the pelvic fin of skates. In the tetrapod lineage, the opposite trend, restricting expression domains of a certain set of *Hox* genes such as the *Hox4* and *Hox5* paralogous groups, might transform the neuromuscular pattern of dual contribution into an exclusive one.

The newly defined expression of *Lbx1* in skate dorsal fins sheds light on the evolutionary origin of appendicular neuromuscular systems (Fig. 5). If paired appendages evolved by a redeployment of the developmental programs in dorsal fins (46), then MMPs with *Lbx1* expression in dorsal fins are likely to be the evolutionary origin of paired appendage muscles. During the origin of paired appendages, the developmental programs of MMPs, including *Pax3* and *Lbx1* expression, might have been redeployed from unpaired to paired appendages, and elaborated by recruiting other signaling pathways such as HGF/MET (47). If paired appendages were assembled *de novo*, then genetic programs of MMPs for paired appendages might have been deployed from MMPs that were already equipped in the development of other muscles such as hypobranchial, dorsal fin, or diaphragm muscles. Further analysis of regulatory regions of *Lbx1* in multiple species including agnathans, which have only unpaired fins, could pursue this hypothesis. Our analysis of neuromuscular development in chondrichthyans illuminates the evolutionary origins and diversification mechanisms of neuromuscular systems of paired appendages.

## Materials and Methods

### Animal husbandry

All processes and protocols for experimental animals were approved by the Institutional Animal Care and Use Committee (IACUC) of Rutgers and University of Chicago. *Leucoraja erinacea* and *Scyliorhinus retifer* embryos were purchased from the Marine Resource Center of The Marine Biological Laboratory. Embryos were fixed by 4% paraformaldehyde (PFA) or Bouin’s solution and subjected to immunostaining, histology, or *in situ* hybridization. Stages were determined by referring to previously published studies (28, 48, 49).

### Characterization of ortholog genes

Cloning and characterization of *Wnt3* and *Fgf8* were described in the previous study(27). *Lbx1, Pax3, Cyp26a1* and *Hox* genes used in this study were cloned from cDNA of stage 23 and 30 skate embryos by reverse-transcriptase PCR and integrated into pCRII-TOPO vector (Invitrogen). PCR cloning primers are listed in Supplementary Table 1. After cloning, each sequence was compared with the previously annotated skate transcriptome (27).

### Whole – mount immunostaining and *in situ* hybridization in chondrichthyans

Whole-mount *in situ* hybridization was performed as previously described (27). The details of the protocol and replicate numbers is in Supplementary Information.

### Paraffin sectioning and HE staining

Two shark and two skate embryos were fixed in Bouin’s solution overnight at room temperature and then washed with 70% ethanol followed by 100% ethanol. Paraffin sectioning of fixed embryos (8 μm) and hematoxylin and eosin staining were performed by the Human Tissue Resource Center at the University of Chicago (https://pathcore.bsd.uchicago.edu/index.php).

### Cryosectioning of skate embryos

After whole-mount *in situ* hybridization, embryos were re-fixed in 4% PFA and immersed in a graded series of sucrose/PBS solutions (10%, 15%, 20%). Embryos were placed in OCT compound (Tissue-Tek) overnight. The following day, embryos were embedded and sectioned (8 μm) by Leica CM3050S.

### Phalloidin staining

Skate embryos were recovered from egg cases and fixed in 4% PFA overnight. The following day, embryos were treated in a sucrose/PBS solution series, embedded, and cryosectioned as described above. Cryosections of four skate embryos were rinsed in PBTriton 3 times and incubated with 10% sheep serum/PBTriton for 30 minutes at room temperature. Sections were washed by PBTriton 3 times and incubated with 1:1000 phalloidin–Alexa488 (Invitrogen) and 1:4000 DAPI in PBTriton for 1 hour. After sections were washed 3 times in PBTriton, fluorescent images were captured on a Zeiss LSM510.

### Reconstruction of innervation in shark embryos

Hematoxylin and eosin-stained sections were photographed with a Leica M205 FCA. Images were imported into Amira 3D reconstruction software (ThermoFisher). Image direction was aligned, and 3D morphology was reconstructed. Nerves innervating the pectoral fin and body wall muscles were segmented manually and pseudo-colored.

### Investigation of *Hox* expression patterns in NT, PAM, and LPM

After whole-mount in situ hybridization, expression domains of *Hox* genes were investigated under a stereomicroscope and photographed (Leica M205 FCA and MC170 HD). To determine the anterior limit of expression of *Hox* genes in NT and LPM, the number of somites lateral to these tissues were counted. Expression at ventral somites was used to identify anterior limits of *Hox* expression in PAM.

### Culture of skate embryos with RA

Five skate embryos were cultured from stage 23–30 in 900 mL of salt water (Instant Ocean) with/without all-trans RA (Sigma; final concentration of 2 × 10^−6^ M). After the culture, embryos were fixed with 4% PFA and immunostained with 3A10 antibody. We repeated this experiment twice, investigating 10 embryos for each negative control and RA treatment.

## Acknowledgements

We thank David Remsen, Scott H. Bennett, and all other members of the Marine Resource Center at the Marine Biological Laboratory for husbandry of skates and sharks. This work was supported by institutional support provided by the Rutgers University School of Arts and Sciences and the Human Genetics Institute of New Jersey; a Marine Biological Laboratory research grant (to T.N.); and the Brinson Foundation and University of Chicago Biological Sciences Division (to N.H.S.). All experiments were approved by the animal committees of Rutgers University (protocol # 201702646) and the University of Chicago (protocol # 71033).

## Author contributions

N.T., D.M., N.A., N.H.S., and T.N. designed the research; N.T., D.M., K.B., K.F., and T.N. performed the research; N.T., D.M., K.B., N.A., and T.N. analyzed data; and N.T., D.M., K.B., G.S., K.F., N.A., N.H.S., and T.N. wrote the paper.

