## Supplementary Information for "The evolutionary origins and diversity of the neuromuscular system of paired appendages in batoids"

### SI Appendix

#### Supplementary Methods

##### Whole–mount immunostaining and *in situ* hybridization in Chondrichthyans

For *Pax3* and *Lbx1* expression, 12 replicates for each gene were investigated. For *Wnt3, Cyp26a1, Fgf8*, and *Hox* genes, 3 embryos for each gene were investigated. The nerve staining method was modified from a previous study (50). Briefly, skate and shark embryos were transferred from salt water to 500 mg/L Tricaine-S and then fixed in 4% PFA overnight. The following day, embryos were washed in phosphate buffered saline (PBS) containing 1% Triton (PBTriton) for 3 hours. Embryos were incubated in 0.25% trypsin/PBS for 5 minutes and immersed in pre-cooled acetone at –20ºC for 10 minutes. After brief rinsing with PBTriton, specimens were placed in blocking solution (PBTriton containing 10% goat serum, 1% dimethyl sulfoxide, and 5% $H_2O_2$) at 4ºC overnight. The following day, the solution was replaced with blocking solution containing 1:50 3A10 antibody for nerve staining (Developmental Studies Hybridoma Bank) or 1:70 myosin heavy chain for muscle staining (Developmental Studies Hybridoma Bank). Embryos were incubated in antibody solution at 4ºC for 72 hours. Embryos were washed with PBTriton for 5 hours and then incubated with blocking solution containing 1:1000 peroxidase-conjugated secondary antibody (Jackson Laboratory, catalogue number 115-035-003). The next day, embryos were washed with PBTriton for 5 hours and subjected to DAB color development. At each stage, we stained nerves of 6 embryos and muscles of 4 embryos.

**Supplementary Figure 1 | The dual contribution of *Pax3* muscle precursor cells into the pectoral fin and body wall muscles.**

*Pax3*-positive cells migrated into the pectoral fin (arrowheads) and body wall muscles (arrows) in skate embryos at stage 29. Scale bar is 1 mm.

**Supplementary Figure 2 | The development of hypaxial muscles in shark embryos at stage 30.**

Ventral view of shark muscles stained by myosin heavy chain antibody at stage 30. Note that pectoral fin muscles (arrow) and body wall muscles (arrowheads) develop at the same axial level. Scale bar is 1 mm.

**Supplementary Figure 3 | *Hoxa9* expression patterns of embryos treated by retinoic acid from stage 23 to 24.**

Whole-mount *in situ* hybridization of *L. erinacea* embryos at stage 23 to 24. (**A**) *Hoxa9* expression in a wild type embryo. LPM expression starts from somite 26 (arrowhead). (B) *Hoxa9* expression in an embryo treated by RA. LPM expression starts from somite 23 (arrowhead). Note that *Hoxa9* expression is shifted anteriorly in the RA+ embryo by 3 somites. Scale bars are 1 mm.

**Supplementary Table 1 | Cloning primers for *Cdx2, Cyp26a1, Lbx1*, *Pax3*, and *Hox* genes of skate**

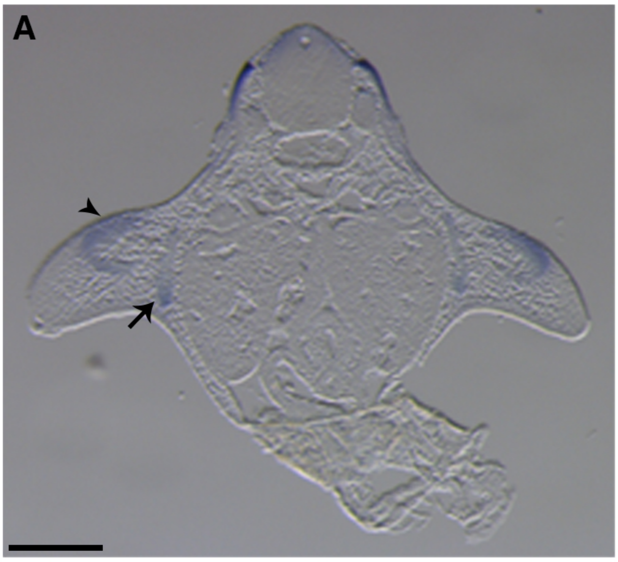


Supplementary Figure 1

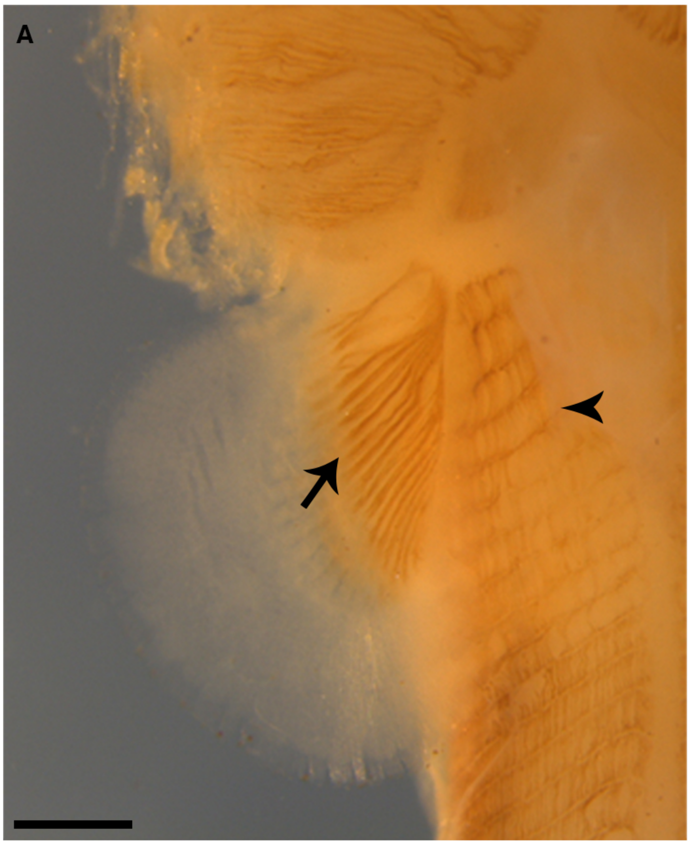


Supplementary Figure 2

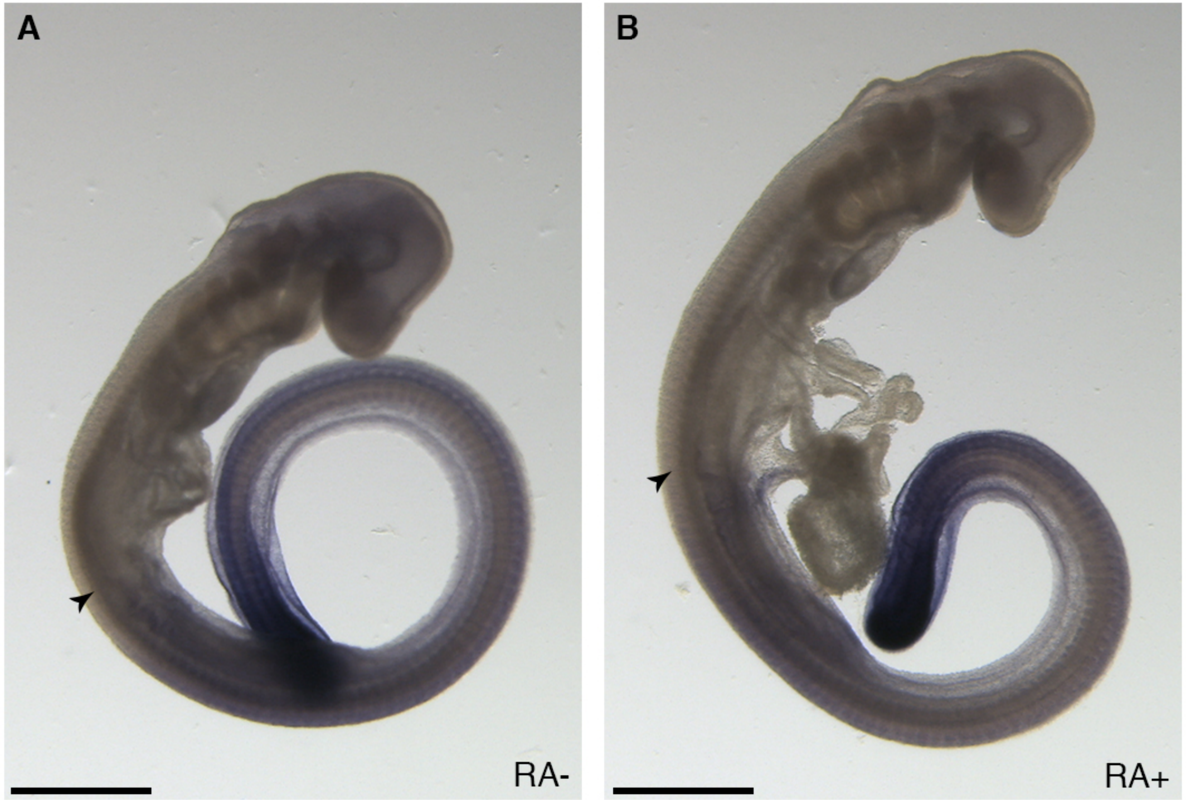


Supplementary Figure 3

Supplementary Table 1

| Gene | Forward Sequence | Reverse Sequence | Length |
|---|---|---|---|
| *Hoxa2* | CAAACCATCCCTAGCCTGAA | TGGACCCATGTATTGCTGAA | 1133bp |
| *Hoxa3* | CTCAATCCAGCACAAGCAAA | GGCTGGAGTCCATGAAAAGA | 1104bp |
| *Hoxa4* | CCCCAATTACACTGGAGGAG | GGCAGTTTGTGATCTTTCTTCC | 209bp |
| *Hoxa5* | TAGACGCACAAACGACCAAG | CAGGCAACAGTACGACCAGA | 1053bp |
| *Hoxa9* | GCTGCCCTTACACCAAACAT | TTTCTTTGCACCACAACACC | 975bp |
| *Hoxa10* | ACCTTATTGCCGTTCTGGTG | TTTTCTCCCCAATTCACTCG | 803bp |
| *Hoxa11* | GTTTAGCTGTGCGAATGCAA | TGCGCTTCTTATCGCCTTAT | 1391bp |
| *Hoxa13* | GAACTTCACCGCAAACCAAT | CGTGCAGGCAGAAATGTAAA | 917bp |
| *Hoxd1* | GTCACCAGGGCTGTGACTTT | AGCGTTCGCTATTTCCACAC | 469bp |
| *Hoxd3* | TCAGACATCCAGTGCCAGAG | TCCCTGAGCTGTGTGATGAG | 1002bp |
| *Hoxd4* | ATCCCAAATTTCCTCCTTGC | GGGTTCCCCTCCAGTGTAAT | 400bp |
| *Hoxd8* | AAGCCAGGAGAAGCCCTAAG | TTCTATCCGGCGGTTGATAC | 929bp |
| *Cdx2* | TTTGTCAGCTTTCGCTCCTT | ATGTACGTGAGYTAYCYT | 623bp |
| *Cyp26a1* | CAATGCTCGACATGGTCTTG | CTTCACAGTGGAGCTGGTCA | 552bp |
| *Lbx1* | CCGTTCAGCATCGAGGATA | GCCTTCATCTCTTCCAGGTC | 473bp |
| *Pax3* | ATGAAGGGTGGGGTGGATA | ATCTGGGACTGAACGTGTCC | 1505bp |
